# *Wolbachia* introgression in Rio de Janeiro remains at sub-optimal levels 30 months after its crash: challenges in the sustainability of *w*Mel interventions for dengue control

**DOI:** 10.1101/2025.07.29.667468

**Authors:** Karine Pedreira Padilha, Gabriela Azambuja Garcia, Rayane Teles-de-Freitas, Franck Jeannot Gnonhoue, Mariana Rocha David, Jessica Corrêa-Antônio, Felipe de Oliveira, Luiz Paulo Brito, Luciana Dias, Ademir Jesus Martins, Vincent Corbel, José Bento Pereira Lima, Gabriel Luz Wallau, Ary Hoffmann, Oswaldo Gonçalves Cruz, Daniel Antunes Maciel Villela, Márcio Galvão Pavan, Rafael Maciel-de-Freitas

**Affiliations:** Laboratório de Mosquitos Transmissores de Hematozoários, Instituto Oswaldo Cruz, Fiocruz, Rio de Janeiro, Brazil; Laboratório de Biologia de Tripanosomatídeos, Instituto Oswaldo Cruz - IOC, Rio de Janeiro, RJ, Brazil; Laboratório de Biologia, Controle e Vigilância de Insetos Vetores, Instituto Oswaldo Cruz, Fiocruz, Rio de Janeiro, Brazil; MIVEGEC, IRD, CNRS, University of Montpellier, Montpellier, France; Departamento de Entomologia e Bioinformática, Instituto Aggeu Magalhães, Fiocruz Pernambuco, Brazil; Department of Entomology and Arbovirology, Bernhard Nocht Institute for Tropical Medicine, Hamburg, Germany; Universidade Federal de Santa Maria, Rio Grande do Sul, Brazil; Pest and Environmental Adaptation Research Group, School of BioSciences, Bio21 Institute, The University of Melbourne, Melbourne, Australia; Programa de Computação Científica, Fiocruz, Rio de Janeiro, Brazil

**Keywords:** *Aedes aegypti*, *Wolbachia*, wMel, arboviruses control, dengue, insecticide resistance, integrative vector management

## Abstract

The deployment of the *Wolbachia w*Mel strain is currently underway in multiple dengue-endemic municipalities across Brazil. The efficacy of this strategy in Rio de Janeiro remains uncertain, primarily due to the difficulty in sustaining high *w*Mel prevalence in regions previously subjected to large-scale releases. A key contributing factor was the routine rotation of insecticides within the framework of Integrated Vector Management (IVM), which led to the use of the larvicide Spinosad for Aedes control in urban areas. This compound was associated with a precipitous decline in *Aedes aegypti* mosquitoes, regardless the *w*Mel infection status. While *w*Mel-uninfected population recovered within weeks, *w*Mel-infected population remained at low levels likely due to the fitness costs imposed by *w*Mel on egg viability. To assess the long-term persistence of wMel following this demographic collapse, we conducted mosquito sampling across 12 neighborhoods in Rio de Janeiro, 30 months after mosquito populations crashed. Our findings reveal that *w*Mel introgression remains suboptimal, with a mean frequency of 9.87% across the sampled areas. Only two neighborhoods exhibited wMel frequencies exceeding 15%, likely reflecting ongoing localized releases. The reduced prevalence underscores the challenges of achieving self-sustaining *w*Mel establishment in complex urban environments and highlight critical considerations for the implementation of *Wolbachia*-based dengue control programs in endemic regions.

**Sponsorship:** This study was supported by grants from the CNPq – Conselho Nacional de Desenvolvimento Científico e Tecnológico (312282/2022-2, 307209/2023-7) and by FAPERJ – Fundação Carlos Chagas Filho de Amparo a Pesquisa no Estado do Rio de Janeiro (E-26/211.159/2019, E-26/204.108/2024, E26/2001.844/2017, E-26/210.335/2022, E-26/210.537/2024).

## Introduction

*Aedes aegypti* (Diptera: Culicidae) is a mosquito species extensively distributed across urban areas of tropical and subtropical regions. It constitutes a major public health concern as the principal vector of several medically important arboviruses, including dengue, Zika, and chikungunya viruses. Among these, DENV is particularly significant due to its widespread global presence and the frequent outbreaks and epidemics it causes, affecting millions of individuals annually.^(1,2)^ The burden of dengue is predominantly concentrated in tropical regions, with Brazil consistently reporting the highest number of cases.^(3)^ In 2024 alone, Brazil registered over 6.6 million suspected dengue cases, marking the highest annual incidence since the virus was reintroduced into the country in the late 1980s – early 1990s.^(4)^

The continued rise in arboviral cases globally, and particularly in Brazil, demonstrates the limited effectiveness of conventional vector control methods, such as chemical insecticides and mechanical interventions.^(5-7)^ This scenario underscores the urgent need for complementary strategies to enhance *Ae. aegypti* population control.^(8,9)^ Among several innovative approaches, the use of the symbiotic bacterium *Wolbachia pipientis* has gained attention. While naturally absent in *Ae. aegypti, Wolbachia* strains have been successfully transinfected from other insect species into *Ae. aegypti*, and some of them have pathogen-blocking properties.^(10-14)^ The spread of *Wolbachia* strains through arthropod populations is favored by a combination of efficient maternal transmission, which is dependent on high *Wolbachia* density in female reproductive systems, and their ability to cause cytoplasmic incompatibility (CI), in which uninfected females are at a disadvantage over infected females because they fail to produce offspring when mated with infected males.^(15)^ It is thereby possible to replace local *Ae. aegypti* populations with high susceptibility to arbovirus with *Wolbachia*-infected equivalents that have a reduced potential for pathogen transmission.

The successful introgression of *Wolbachia* into native *Ae. aegypti* populations involves reaching a high prevalence in the target area with long-term stability. Monitoring efforts are essential to evaluate the proportion of mosquitoes carrying *Wolbachia*, thereby assessing the effectiveness and persistence of the introgression. Notably, *Wolbachia* prevalence in *Ae. aegypti* populations can fluctuate over time and often displays a markedly heterogeneous spatial distribution.^(16-20)^ These fluctuations may be driven by several factors, including reduced fitness of *Wolbachia*-infected individuals, insecticide resistance, migration of wild-type mosquitoes, and environmental conditions such as high temperatures, which can particularly affect the stability of the *w*Mel strain of *Wolbachia*. ^(21-31)^

In the last ten years, *Wolbachia* deployments have been conducted in several cities worldwide from countries as diverse as Australia, Brazil, Colombia, Indonesia, Malaysia, Singapore, and Vietnam, with reports of epidemiological impact.^(16,32-39)^ An intriguing pattern can be observed in Brazil: the neighboring cities of Rio de Janeiro and Niteroi, although separated each other by a bridge, seems to present a very different introgression profile and epidemiological outcome. In Niterói, *w*Mel mosquitoes were deployed to the full extension of the city limits, which the authors reported to be associated with a substantial reduction of 69% of dengue incidence rate compared to predefined control areas.^(40)^ The outcomes in Rio de Janeiro were more modest. Despite releasing 67 million *w*Mel *Ae. aegypti* mosquitoes during 2017-2019, an average of only 32% *w*Mel introgression level into the wild population was achieved, which the authors reported to be associated with a reduction of 38% and 10% in dengue and chikungunya notifications, respectively.^(41)^ Although *w*Mel introgression in Rio de Janeiro remained stable at 30-40% frequency after releases ended, in June 2022 a significant decline in *w*Mel frequency was observed.^(42)^ The drop in *w*Mel frequency was associated with the introduction of the larvicide spinosad (Natular™, Clarke Mosquito Control Products, St. Charles, USA), which disrupted both *w*Mel-infected and - uninfected populations but only the latter was able to recover after an initial population crash likely due to fitness costs of *w*Mel on *Ae. aegypti* mosquitoes.^(29,30,43)^ In this article, we aim to update *Wolbachia* introgression in Rio de Janeiro, 30 months after *Ae aegypti* population crashed in the city, and 20 months after the last observation.^(42)^ By doing so, we plan to provide a snapshot of the current bacteria introgression status in Rio de Janeiro, shedding light on the long-term sustainability of this strategy to mitigate dengue transmission in endemic settings.

## Materials and Methods

### Study area

Field collections of adult mosquitoes were conducted in the same neighborhoods of Rio de Janeiro as in the longitudinal sampling performed between August 2021 and March 2023.^(42)^. The total area covers 12 urban neighborhoods: six of them received previous *w*Mel deployment between 2017-2019 from the World Mosquito Program (hereafter named as neighborhoods A-F) and other six did not (named as G-L) (Figure 1). This region covers an area of 33.5 km^2^ (38.6% of the WMP release area) where approximately 495,000 people live (55.6% of the population under *w*Mel intervention), encompassing some of the most neglected communities in Rio due to urban violence.

**Figure 1.**
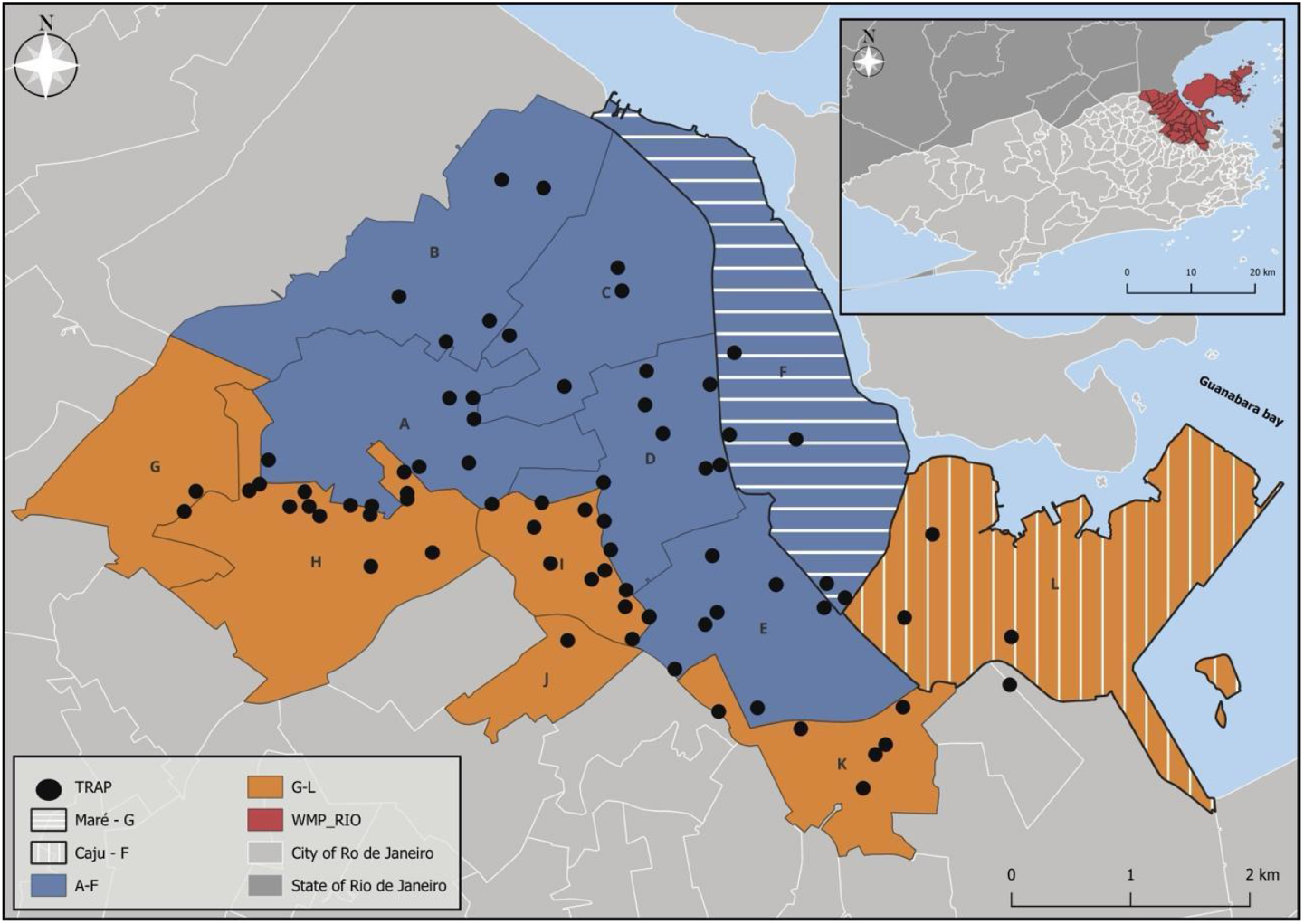
Map of the city of Rio de Janeiro showing in red the total area where mosquito releases were conducted by the World Mosquito Program (WMP). Neighborhoods included in the study are highlighted in blue (release areas; A: Complexo do Alemão; B: Olaria; C: Ramos; D: Bonsucesso; E: Manguinhos; F: Maré) and orange (non-release areas: G: Engenho da Rainha; H: Inhaúma; I: Higienópolis; J: Maria da Graça; K: Benfica; L: Caju). BG-Sentinel trap locations are marked with black dots. The neighborhoods of Maré and Caju are indicated with horizontal and vertical hatching, respectively.

### Mosquito sampling

Between November-December 2024, we installed 75 BG-Sentinel traps in the same 12 neighborhoods and dwellings sampled in Pavan et al. (2025).^(42)^ We investigated whether any changes in *w*Mel frequency occurred 30 months after the crash in *Ae. aegypti* populations was observed. The traps were visited weekly until a predefined number of five *Ae. aegypti* females were obtained. Captured mosquitoes were morphologically identified through taxonomic keys^(44)^ and individually screened for the presence of *Wolbachia w*Mel and relative density.

### In silico design and validation of oligonucleotides (primers) sets

*Wolbachia* density was estimated through qPCR using specific primers that were designed in this study for SYBR Green-based real-time PCR and amplify 115-bp of the *Wolbachia* surface protein gene (*wsp*; GenBank gene ID CP090948) of *w*Mel and 106-bp of the *Ae. aegypti* ribosomal protein S6 gene (*rps6*; GenBank gene ID 5563590). Briefly, genomic sequences were obtained from the Bacterial and Viral Bioinformatics Resource Center v. 3.49 (https://www.bv-brc.org/) and NCBI (https://ncbi.nlm.nih.gov/) databases and sequences were aligned with the MultAlin tool available at (http://multalin.toulouse.inra.fr/multalin/. We used *wsp* gene sequences of different *Wolbachia* strains (*w*Mel, *w*AlbA and *w*AlbB) aiming to select a region that would allow specific amplification of the *w*Mel strain. In the case of the *rps* gene, primers were designed in this highly conserved region of *Ae. aegypti*. Primer design was conducted in silico using an online tool (https://primer3.ut.ee/) to generate nucleotide pairs flanking target regions ranging from 50 to 150 base pairs. Finally, the quality of the designed oligonucleotides was evaluated using an online platform specifically developed for this purpose (https://www.idtdna.com). Primers were designed based on key thermodynamic parameters to ensure the specificity and efficiency of qPCR amplification. We selected primers with melting temperatures (Tm) above 45 °C and the difference between the Tm of forward and reverse primers lower than 5 °C. Additionally, the Gibbs free energy (ΔG) was calculated to assess the propensity for the formation of stable secondary structures, which the selected nucleotides had ΔG > -3 kcal/mol and ΔG > -10 kcal/mol to form hairpins and dimers.

The primers originally described by Ross et al. (2017)^(45)^ for *rps* presented a very low ΔG, favoring the formation of dimers. The new primers for *rps* are entirely located in the third exon of the *RpS6* gene, ensuring that amplification occurs exclusively in the transcribed regions, increasing efficiency and specificity. Comparative analyses indicate that the *RpS6* gene is highly conserved between *Ae. aegypti* and *Ae. albopictus*, which suggests that the same primers may be applicable to both species. Regarding the primer pairs for *w*Mel, the forward primer was designed in a region highly conserved among the *w*Mel, *w*AlbA and *w*AlbB strains, presenting only three specific nucleotides that differentiate wMel from *w*AlbA. On the other hand, the reverse primer demonstrates greater specificity for wMel, containing ten nucleotides unique to this strain and eight nucleotides shared between *w*Mel and *w*AlbA. Detailed information regarding the designed oligonucleotides is provided in Table 1.

**Table 1:**
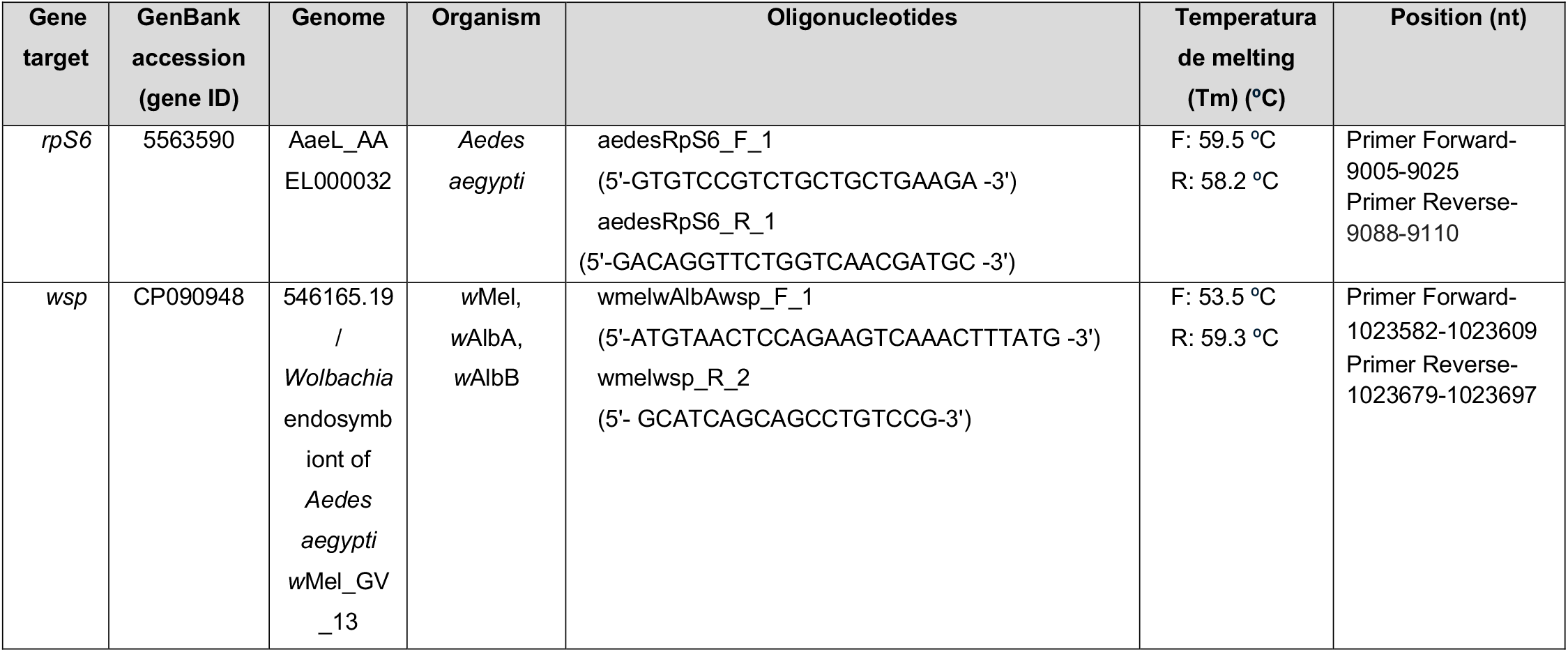
Overview of the specific oligonucleotides data. Details of designed oligonucleotides, including their names, accession IDs, genomic positions, target organisms, and nucleotide sequences. In the table: In grey: exon; green : regions conserved among *w*Mel, *w*AlbA, and *w*AlbB; in blue: regions shared between *w*Mel and *w*AlbA; in yellow: regions unique to *w*Mel.

Laboratory-reared *Ae. aegypti* mosquitoes infected with *w*Mel (F5 generation from Tubiacanga, Rio de Janeiro, Brazil) and not infected with *w*Mel (Paea strain) were used to test primer efficiency and specificity. Mosquitoes were reared in an incubator (Solab model SL-200/364) from egg to adult stage, under a photoperiod of 12 h of light and dark (LD 12:12) and at 28°C. Females aged 0 to 3 days were flash-frozen in liquid nitrogen. Subsequently, each individual body was homogenized using a pestle, and the resulting material was stored in 350⎰l of Buffer RLT Plus (reagent of the allprep DNA/RNA Mini Kit extraction kit, Qiagen, Hilden, Germany) in a -80ºC freezer until subsequent extraction of genetic material. Total DNA and RNA were extracted using the Allprep DNA/RNA Mini Kit (Qiagen, Hilden, Germany), according to the manufacturer’s protocol and the DNA was used in the primer testing assays. The concentration of the extracted DNA was determined using the Qubit 4 Fluorometer quantification system (Thermo Fisher Scientific, Waltham, USA). A pool of ten individuals (each individual sample initially had ∼10 ng/µl of DNA) of each positive (*w*Mel-infected) and negative (*Wolbachia*-free mosquitoes) control was used to calculate oligonucleotide efficiency and specificity through serial dilutions ranging from 1/1x to 1/32x (12.5-400-ng total DNA). The experiments were performed in a QuantStudio 6, using 7.5 µl of Power SYBR Green PCR Master Mix (Thermo Fisher Scientific), with 0.75 µl of each oligonucleotide at 10 mM, 4 µl of DNA, and 2 µl of water in the QuantStudio™ 6 Flex Real-Time PCR System (Applied Biosystems, Waltham, MA, USA), under the following regime: an initial incubation at 50 °C for 2 minutes, followed by an initial denaturation at 95 °C for 10 minutes. The amplification protocol consisted of 40 cycles of denaturation at 95 °C for 15 seconds and annealing/extension at 60 °C for 1 minute. A melt curve analysis was subsequently performed under the following conditions: 95 °C for 15 seconds, 60 °C for 1 minute, and a gradual increase to 95 °C for 15 seconds. All reactions were done with three independent replicates.

### DNA extraction and *Wolbachia* quantification in field sampled mosquitoes

Individual females of *Ae. aegypti* were massed in L15 culture medium (Leibovitz) in a Tissue Lyser II apparatus (Qiagen, Hilden, Germany) for 60 s at 50 Hz and the DNA was extracted using the NucleoSpin®Tissue extraction kit (Macherey-Nagel, Düren, Germany) following the manufacturer’s protocol. The DNA was quantified in a Qubit 4 Fluorometer quantification system (Thermo Fisher Scientific, Waltham, USA). The relative quantification of *w*Mel in whole mosquitoes was performed with quantitative PCR by amplifying the *Wolbachia wsp* gene, using the *rps6* of *Ae. aegypti* as an endogenous control.^(46)^ Quantitative PCR was performed following the same parameters, reagents, equipment and were run under the same cycling conditions as previously outlined for the primers test. The highest resulting delta threshold cycle (ΔCt) was used as a reference sample for delta delta Ct calculations (ΔΔCt).^(47)^

### Statistical analysis

The relative frequency of wMel was compared after the classification of neighborhoods according to the presence/absence of previous *Wolbachia* deployments. We fitted a generalized linear model (GLM) with a negative binomial distribution with the number of *Wolbachia*-infected mosquitoes per trap as the depended variable and the occurrence of releases as the explanatory variable. In addition, we included a log-transformed offset of the total number of mosquitoes tested pert trap, allowing for the interpretation of the model coefficients in terms of relative rates. Model fit was examined by checking heteroscedasticity, residuals dispersion and the presence of outliers with the R package DHARMa.^(48)^ To test the significance of the predictor variable, we used the type II analysis of deviance. These analyses were performed in the R environment (version 3.6.2).^(49)^

Assuming *Wolbachia*-mediated arbovirus blocking effect is often density dependent^(50)^ we also quantified the endosymbiont density relatively to the *rps6* of *Ae. aegypti* in field-caught mosquitoes. The *w*Mel density was compared between females captured in areas that received or not *Wolbachia* deployments, as well as with a Fiocruz lab colony using the Kruskal-Wallis test, followed by Dunn multiple comparisons post hoc tests.

## Results

### Primer evaluation

Melting curve analysis revealed a single peak for the *wsp* gene, suggesting that only one fragment is amplified per reaction (Figure 2). The reaction efficiency for the oligonucleotides targeting the *wsp* gene of *w*Mel falls within the optimal range, with 92.3% efficiency. Regarding the *Ae. aegypti rps6* gene, amplification occurred in both *w*Mel-infected and *Wolbachia*-free mosquitoes (Figure 3), and the melting curve also presented a single peak in both mosquito groups. The efficiency of the primers was 103.0%, also falling into the optimal range.

**Figure 2:**
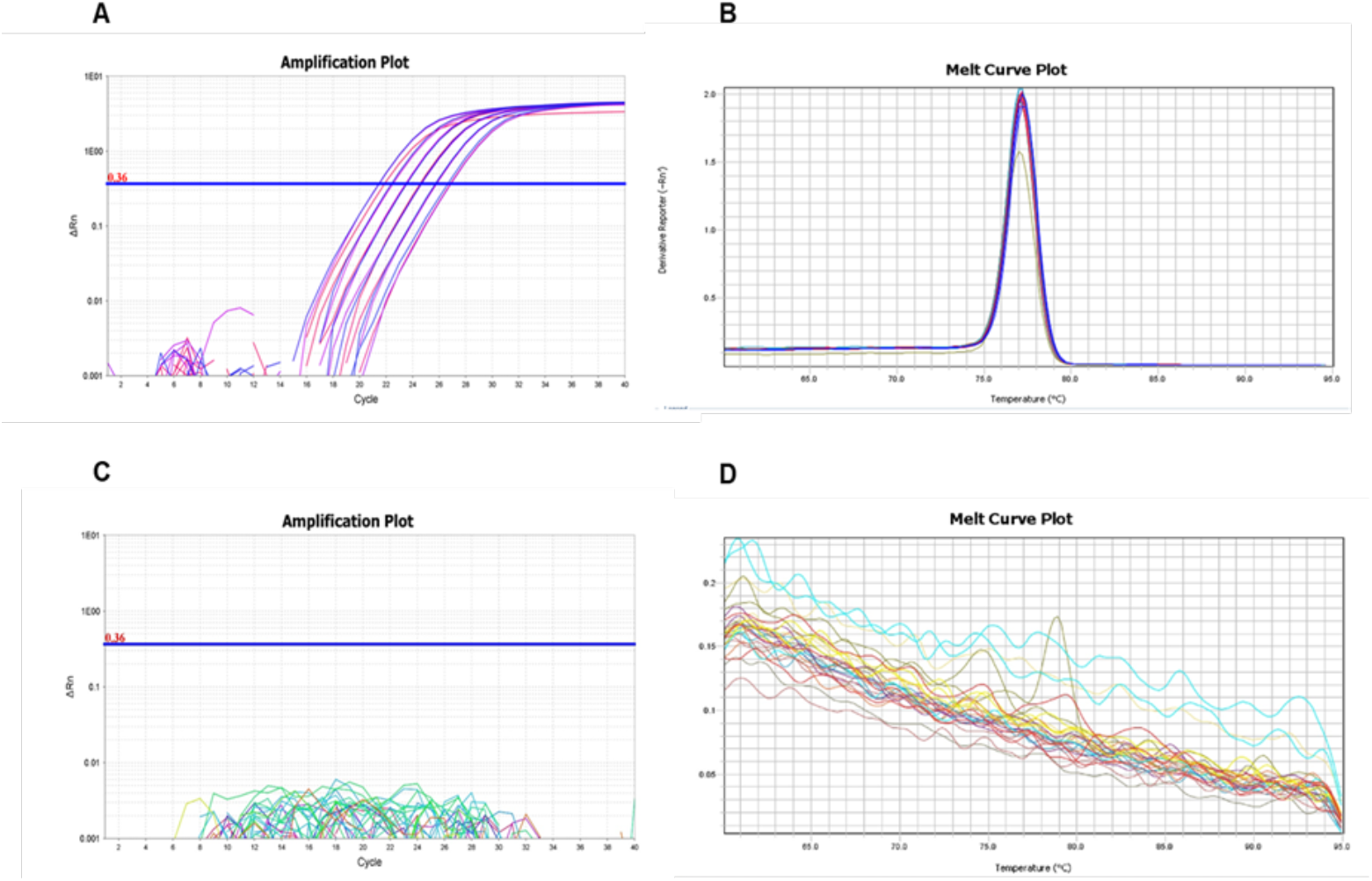
Oligonucleotides for the wsp gene of *w*Mel – A) Amplification curve for the wsp gene of *Wolbachia w*Mel, using DNA from *Ae. aegypti* females with *w*Mel; B) Melting curve for the wsp gene of *Wolbachia w*Mel, using DNA from *Ae. aegypti* females infected with *w*Mel; C) Amplification curve for the *wsp* gene of *w*Mel, using DNA from *Wolbachia*-free *Ae. aegypti* females (*PAEA* strain); D) Melting curve for the *wsp* gene of *Wolbachia w*Mel, using DNA from *Wolbachia*-free *Ae. aegypti* females (*PAEA* strain).

**Figure 3:**
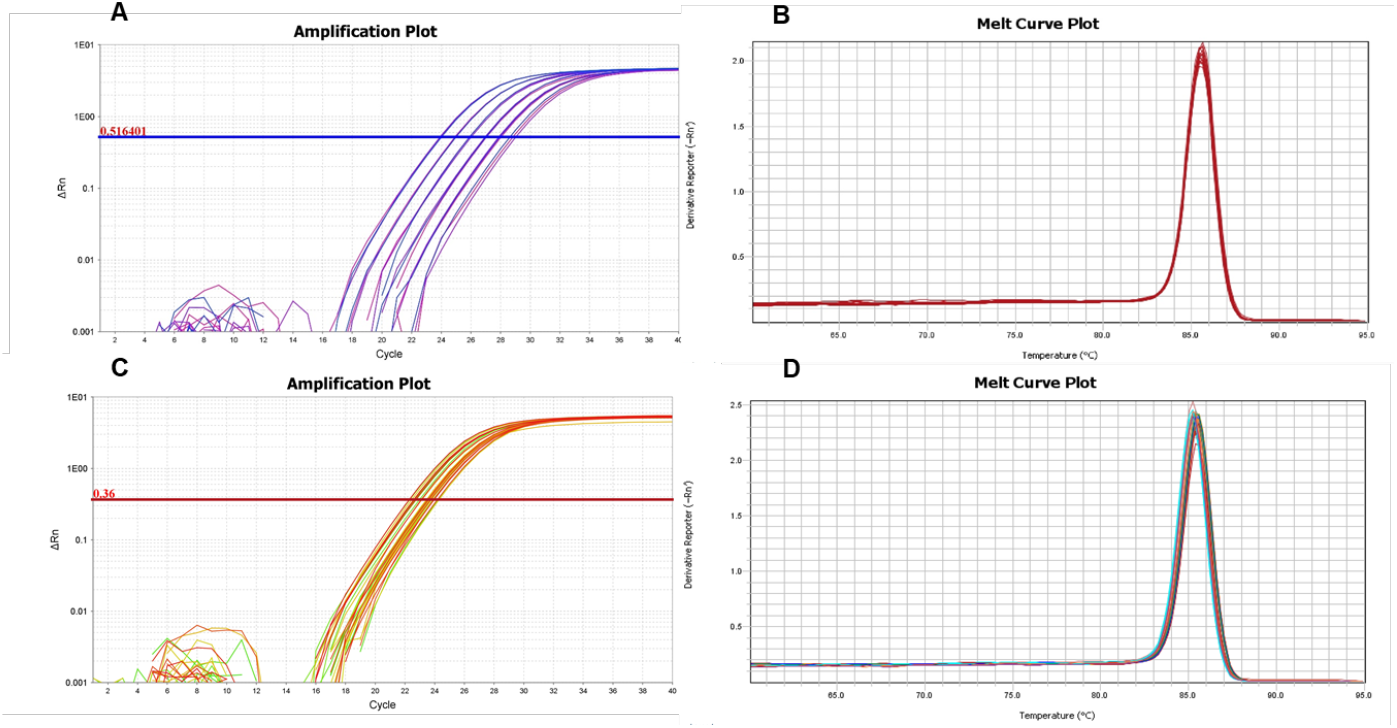
Oligonucleotides for the ribosomal protein S6 (rps6) gene of *Aedes aegypti* – A) Amplification curve for the *rps6* gene of *Wolbachia w*Mel, using DNA from *Ae. aegypti* females infected with *w*Mel; B) Melting curve for the *rps6* gene of *Wolbachia w*Mel, using DNA from *Ae. aegypti* females infected with *w*Mel; C and D) Amplification and curves for the *Aedes rps6* gene, using DNA from 10 individualized *Ae. aegypti* females without *Wolbachia* (Paea lineage).

### Frequency of *Wolbachia* wMel in field *Aedes aegypti*

In total, we tested 375 *Ae. aegypti* females from 12 neighborhoods over the period in which traps remained in the field. There were no significant differences on the number of *Wolbachia*-infected mosquitoes according to the occurrence of previous *Wolbachia* deployments. The coefficient for this explanatory variable was negative but not statistically significant (estimate = –0.3514, z = –0.752, p = 0.452). This finding was supported by the Type II analysis of deviance (χ^2^ = 0.568, df = 1, p = 0.451), indicating a similar frequency of *w*Mel mosquitoes in areas adjacent to release zones (Figure 4), as previously reported (Pavan et al. 2025).

**Figure 4:**
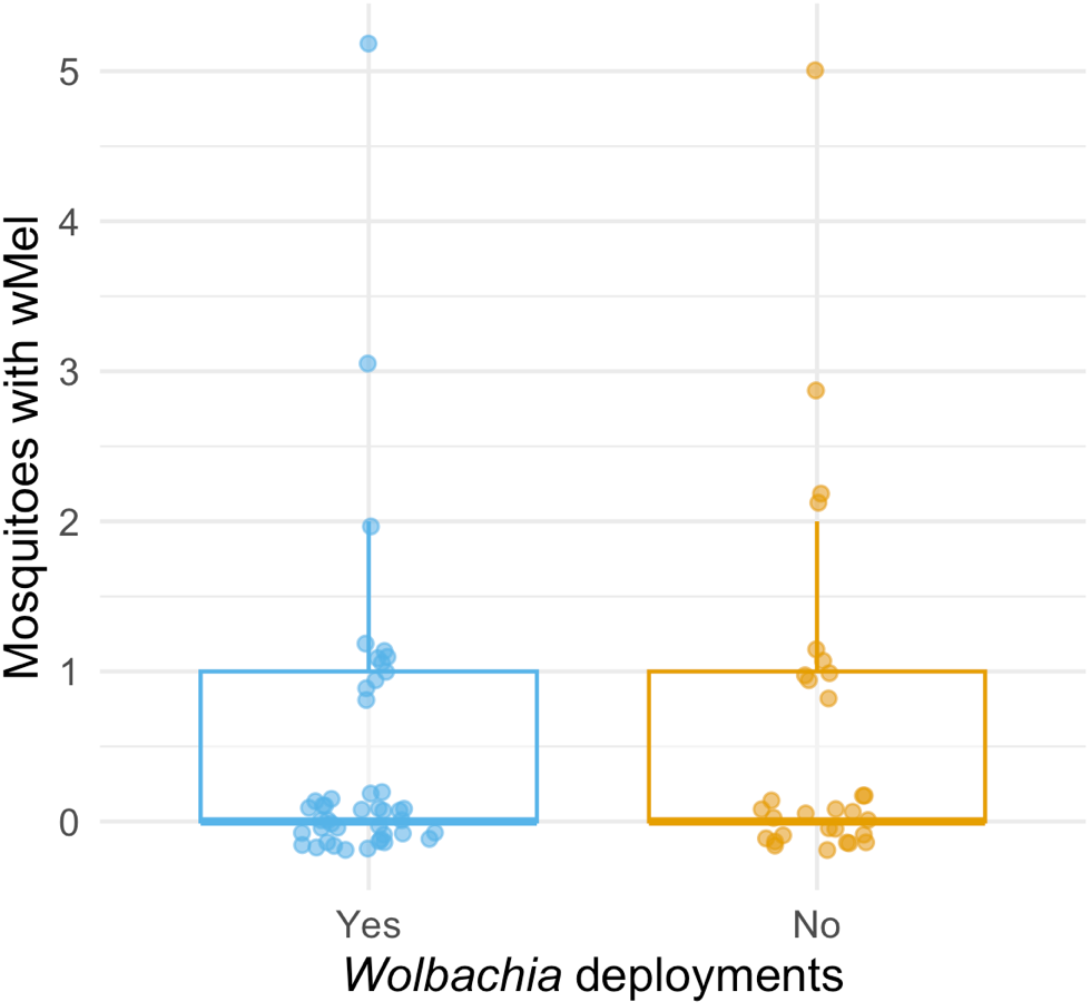
Frequency of mosquitoes (out of 5) detected with the wMel strain of *Wolbachia* per trap in neighborhoods with and without previous mosquito releases by the WMP.

The frequency of *w*Mel in release and non-release areas did not change much between the period of the population crash and the subsequent collection. In March 2023, the *Wolbachia* frequencies in release and non-released areas were 13.2% and 5.01%, respectively. In November 2024, the frequency of *w*Mel in released areas dropped to 8%, whereas in the non-release areas it increased to 12% (Figure 5A). Additional information regarding *w*Mel frequency is available if we aggregate data by neighborhood. Considering the neighborhoods with previous releases (A-F), the *w*Mel frequency was reduced between March 2003 and November 2024, with the exception of the area F. Regarding the neighborhoods in which no *w*Mel deployment was carried out, in the areas G and J no *Wolbachia*-infected mosquitoes were captured in both evaluation periods. However, a significant increase in *w*Mel frequency was observed in areas K and L, both located in the southern limit of our study area (Figure 5B). Remarkably, four neighborhoods had no *w*Mel mosquitoes captured in the BG-Sentinel traps by the end of March 2023. Of these, one had previous releases (A), and three others were without previous deployments (G, H, J). On November 2024, the A and H sites had a frequency of 1.54% and 4.44% of *w*Mel-infected mosquitoes, respectively, whereas sites G and J remained without *w*Mel-infected mosquitoes (Figure 5B).

**Figure 5:**
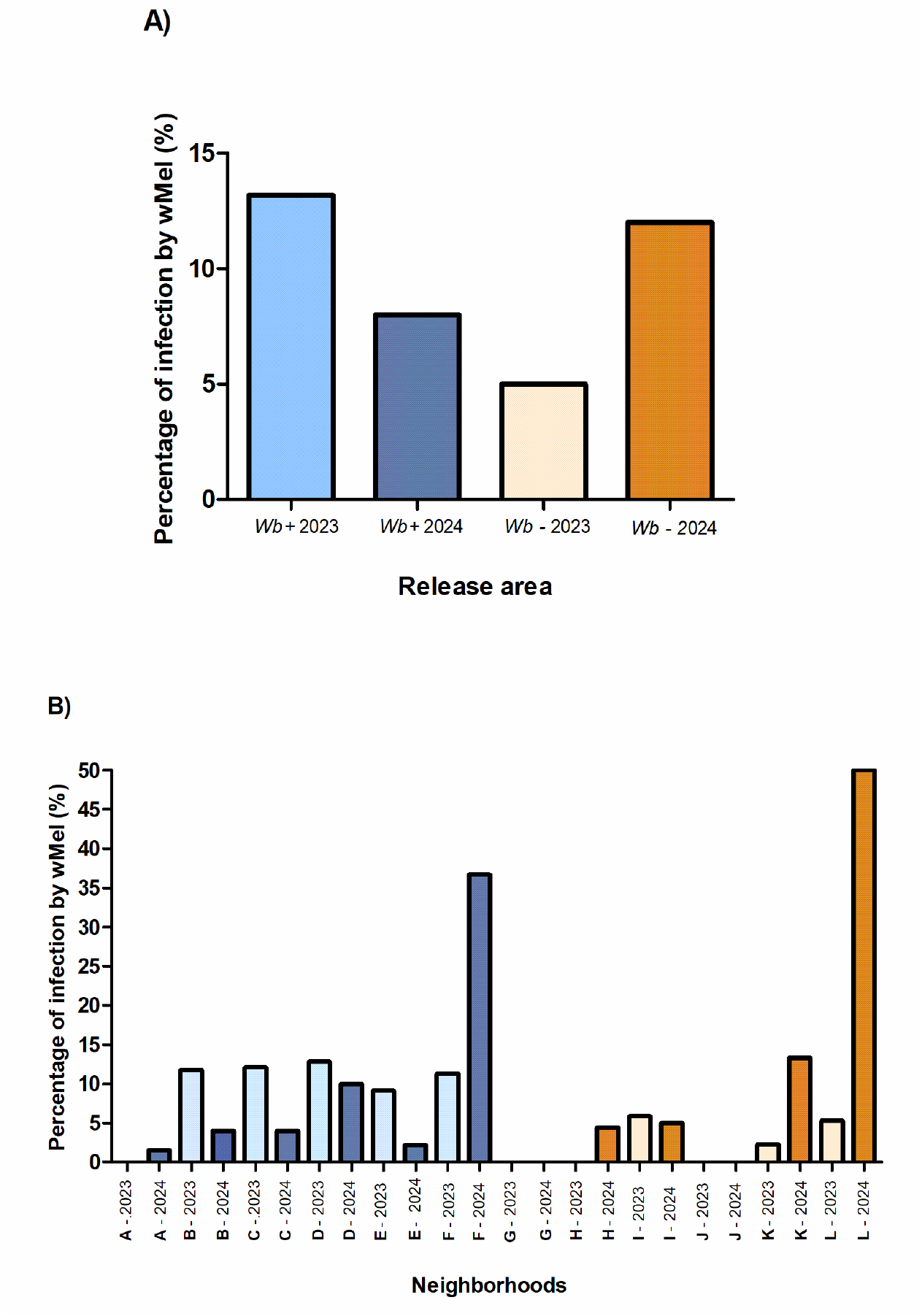
Frequency of *Wolbachia* in female *Aedes aegypti* from the field, using the raw data from Pavan et al. (2025) collected between Feb-Mar 2023 and Nov 2024. In light blue (2023 data) and blue (2024 data) are the average frequencies of *w*Mel in neighborhoods which received *Wolbachia* deployments; in light orange (2023 data) and orange (2024 data) are the frequencies of *w*Mel in areas with no previous *Wolbachia* releases. A) *w*Mel frequency of neighborhoods with and without previous releases. B) Data presented per neighborhood. Notably, Engenho da Rainha and Maria da Graça had no *w*Mel-infected *Ae. aegypti* captured in both periods.

The density of the *w*Mel strain in *Ae. aegypti* mosquitoes was consistent between those individuals caught in areas with or without previous deployment, but those from the lab colony had a higher *Wolbachia* density than the ones collected in neighborhoods that had no releases (H= 11.23, df= 2, p=0.003, followed by Dunn: Z= 18.28, p<0.05)(Figure 6).

**Figure 6:**
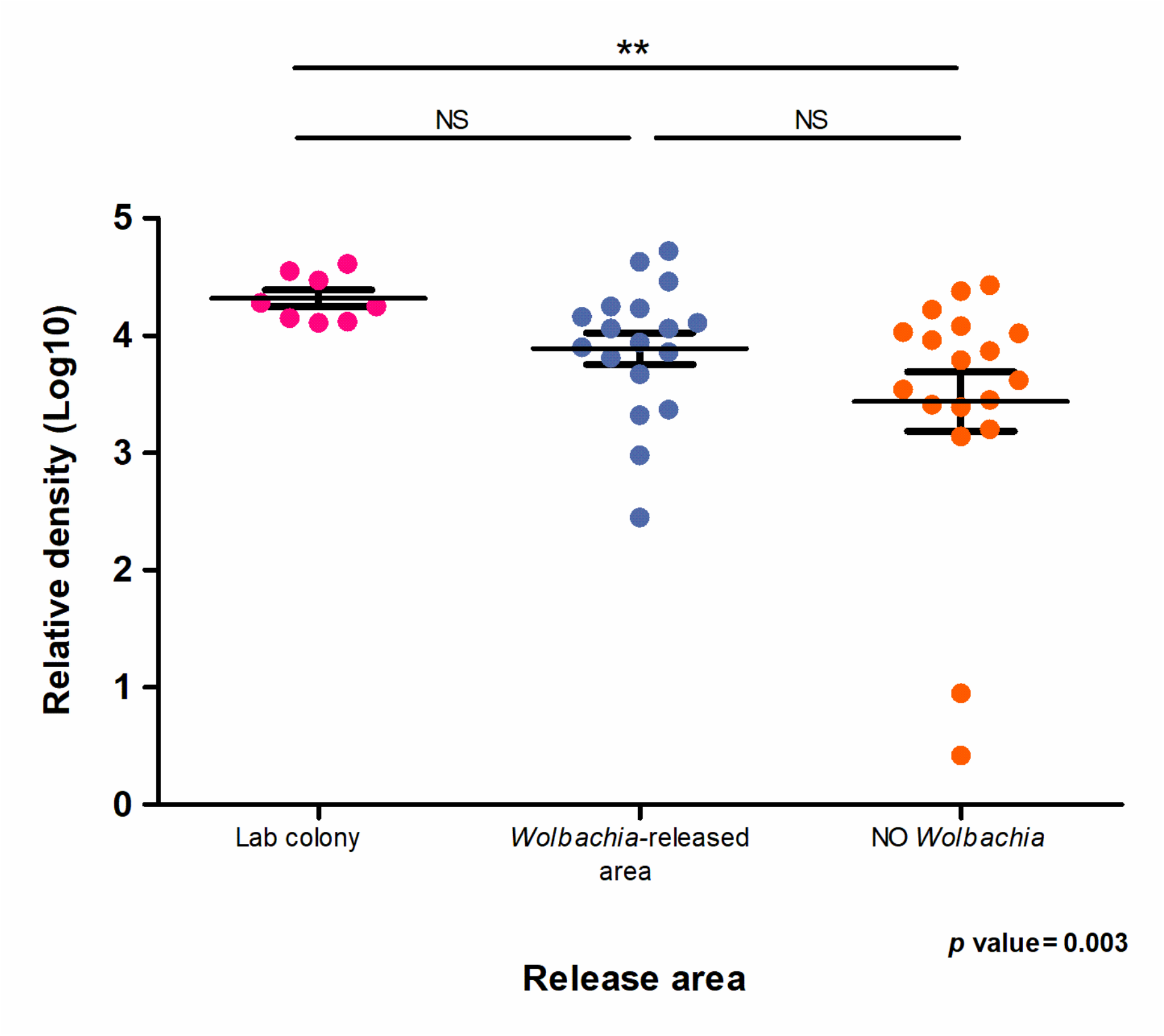
Relative density of *Wolbachia* in *Aedes aegypti* females from the *w*Mel colony and from the field, captured in BG-Sentinel traps installed in areas with or without previous *Wolbachia* releases. The *wsp* gene was used as a target and the *rps6* gene as an endogenous normalizer. Each symbol represents one single mosquito female.

## Discussion

The deployment of *Aedes aegypti* carrying *Wolbachia pipientis* to mitigate dengue transmission has been underway in different epidemiological settings. In Australia, the *w*Mel strain of *Wolbachia* successfully established in local *Ae. aegypti* populations, reaching fixation and maintaining high prevalence accompanied by a 96% reduction in dengue incidence.^(18,51)^ Similarly, in Yogyakarta, areas achieving >70% *Wolbachia* prevalence reported a 77% reduction in dengue cases and an 86.2% decline in dengue-related hospitalizations.^(52)^ The deployment of *w*AlbB in Malaysia resulted in an initial average 40% reduction of dengue cases in the intervention sites, rising to > 60% afterwards.^(34,35,53)^ The release of *w*Mel strain in Brazil had divergent patterns in the neighboring cities of Rio de Janeiro and Niterói for reasons that remain unclear. In Niterói, despite heterogeneous introgression patterns across landscapes, an overall 69% reduction in dengue transmission was obtained^(40)^, while in Rio de Janeiro, a fluctuating *w*Mel prevalence around 32% led to modest reductions in dengue (38%) and chikungunya (10%) incidence.^(41)^ Considering the epidemiological outcome of *Wolbachia* seems to be tightly connected with the level of introgression achieved and the maintenance of high frequency over time, the spatiotemporal evaluation of *Wolbachia* is of utmost relevance to determine its success.

The World Mosquito Program released 67 million *Ae. aegypti* mosquitoes to cover approximately 6.9% of Rio de Janeiro city’s area between August 2017 and December 2019.^(41)^ From August 2021 to March 2023 an independent monitoring was able to follow significant changes in *w*Mel frequency.^(42)^. The *w*Mel frequency remained relatively stable between 30–40% until June 2022, when a sharp decline in both *w*Mel-infected and uninfected *Ae. aegypti* populations was observed in Rio de Janeiro. This coincided with the introduction of the new larvicide spinosad, to which mosquitoes showed full susceptibility. Most likely, the fitness cost associated with this *Wolbachia* strain on mosquito life-history traits prevented its recovery after the spinosad application coverage was reduced by the local public health office.^(28,30,31,42,55-57)^ Our monitoring revealed that in March 2023 only ∼10% of field-caught mosquitoes had *w*Mel. In the present study, we investigated the current profile of *w*Mel introgression in Rio de Janeiro to evaluate potential spatiotemporal changes in *Wolbachia* dynamics. *Wolbachia* frequency remained at low levels similarly to those observed in Pavan et al. (2025)^(42)^. The persistence of *w*Mel frequency at ∼10% suggest its stabilization over the last couple of years in Rio de Janeiro. Considering the reproductive advantage conferred by *Wolbachia* is frequency-dependent, its persistence at the low frequencies reported herein will likely be insufficient to trigger a further increase without needing further releases and, importantly, be ineffective for protecting against dengue outbreaks.^(15,58)^

The introduction of the *w*Mel strain of *Wolbachia* into native *Aedes aegypti* populations has been proposed as a promising and sustainable strategy for arbovirus control, based on the assumption that, once a high prevalence threshold is surpassed, *Wolbachia* frequencies remain stable over time.^(59-61)^ However, the long-term persistence of *Wolbachia* in field populations appears to be highly dependent on a range of ecological and anthropogenic pressures, including insecticide exposure and climatic variability. In Brazil, the initial field release of *wMel*-infected *Ae. aegypti* occurred in the geographically isolated fishing community of Tubiacanga. Although early invasion dynamics were promising, *Wolbachia* frequencies declined sharply following the cessation of releases, coinciding with intensive household level applications of pyrethroid-based adulticides.^(16,62)^ Subsequently, a more extensive collapse in *wMel* prevalence was observed across a broader urban area, temporally associated with the expanded use of the larvicide Spinosad.^(41)^ Similar patterns of instability have been documented elsewhere; for example, in Tri Nguyen village, Vietnam, *wMel* frequencies, which had surpassed 90% during cooler periods, fell to approximately 24% during the hot-dry season, suggesting a pronounced sensitivity to seasonal thermal stress.^(20)^ In Rio de Janeiro, seasonal oscillations in *Wolbachia* prevalence have likewise been recorded, with the lowest frequencies observed during the summer months—coinciding with the peak incidence of dengue transmission.^(41)^ As in Tri Nguyen village, results in Rio de Janeiro also showed small-scale heterogeneity, but most likely due to additional releases conducted by the WMP.^(63)^ The observation that only the neighborhoods of Maré (área F) and Caju (area L) exhibited an increase in *w*Mel frequency is likely attributable to successive rounds of mosquito releases, as reported in local media sources.^(63)^ The necessity of supplementary releases in areas where *Wolbachia* frequencies remain below optimal thresholds should be considered an integral and ongoing component of vector control programs, particularly in light of the environmental factors that can influence *w*Mel dynamics. Given the historical role of Rio de Janeiro as a major port of entry for both dengue serotypes and genotypes, ensuring that *w*Mel frequencies remain at protective levels in previously targeted areas should take precedence over the expansion of releases into new localities.^(64,65)^

An additional consequence of suboptimal *w*Mel frequencies in the field is the reduced level of community protection, as lower prevalence of *Wolbachia* is expected to result in diminished interruption of arbovirus transmission. Notably, in 2024, Rio de Janeiro experienced its most severe dengue outbreak in the past decade.^(66)^ This increase is not directly related to the drop in *Wolbachia* frequency given that prior releases conducted by the WMP covered less than 7% of the city’s total area, but it does emphasize the challenges in disease control moving forward. The present study focuses on approximately 38.6% of the geographic region where *w*Mel-infected mosquitoes were deployed. Given this limited spatial coverage, the releases are unlikely to have exerted a substantial impact on citywide transmission dynamics. Nonetheless, these findings underscore the importance of prioritizing the reinforcement of *Wolbachia* interventions in areas with previous but suboptimal introgression, rather than expanding efforts into new localities before establishing stable and protective *wMel* prevalence in existing ones.

The *w*Mel density of field-caught mosquitoes from the area without previous releases was significantly lower than the lab colony, following the same pattern as observed before.^(43)^ As extensively reported, high temperatures can negatively impact *w*Mel density in *Ae. aegypti*.^(23,26,27)^ Considering lab mosquitoes are reared in optimum insectary conditions, with a stable temperature of 27 ± 2°C, *Wolbachia* density of lab colony mosquitoes is expected to be high. Regarding arbovirus mitigation, it is well known that the pathogen-blocking phenotype associated with *Wolbachia* depends on its density.^(67-69)^ Therefore, seasonal fluctuations in *w*Mel density, especially a decrease in the warmer periods in which dengue transmission is more intense, may affect *w*Mel effectiveness. In fact, preliminary evidence highlighted the presence of *w*Mel not reduced dengue transmission in Rio de Janeiro during the 2024/2025 season.^(70)^

This study has limitations. The sampling strategy encompassed only a subset of the *w*Mel deployment area, thereby constraining the ability to generalize findings to non-monitored regions. Additionally, BG-Sentinel traps were preferentially installed in the same households as those used in Pavan et al. (2025)^(42)^, which facilitates temporal comparison but may perpetuate any sampling biases present in the original dataset. Despite these constraints, the study offers a robust and independent assessment of the spatiotemporal distribution of the *wMel* strain in Rio de Janeiro. These findings highlight the critical importance of adopting an Integrated Vector Management (IVM) framework to enhance the efficiency and sustainability of vector control efforts and disease prevention strategies and to engage in ongoing monitoring and adaptative releases when deploying *Wolbachia*.

## Acknowledgements

We would like to thank Marcelo Celestino Santos and Mauro Menezes Muniz for conducting the field work. This study was supported by grants from the CNPq – Conselho Nacional de Desenvolvimento Científico e Tecnológico (312282/2022-2) and by FAPERJ – Fundação Carlos Chagas Filho de Amparo a Pesquisa no Estado do Rio de Janeiro (E-26/211.159/2019, E-26/204.108/2024, E26/2001.844/2017, E-26/210.335/2022, E-26/210.537/2024).

## Author’s contribution

Conceptualization (MGP, FJG, GAG, OGC, DV, RMF), Data curation (FJG, MRD, FO, LPB, LD), Formal analysis (MRD, JCA, KPP, GAG, FO, OGC, DV), Investigation (MGP, GAG, AJM, VC, JBPL, GLW, AH, OGC, DV, RMF), Methodology (MGP, FJG, JCA, KPP, FO, LPB, LD, OGC, DV), Project administration (MGP, DV, RMF), Resources (MGP, DV, RMF), Supervision (MGP, AJM, JBPL, AH, DV, RMF), Visualization (JCA, KPP, LPB, LD, OGC, DV), Writing -original draft (MRD, MGP, OGC, DV, RMF), Writing – review & editing (MGP, FJG, MRD, JCA, KPP, GAG, FO, LPB, LD, AJM, VC, JBPL, GLW, AH, OGC, DV, RMF).

